# Population structure and inbreeding in wild house mice (*Mus musculus*) at different geographic scales

**DOI:** 10.1101/2022.02.17.478179

**Authors:** Andrew P Morgan, Jonathan J Hughes, John P Didion, Wesley J Jolley, Karl J Campbell, David W Threadgill, Francois Bonhomme, Jeremy B Searle, Fernando Pardo-Manuel de Villena

**Affiliations:** Department of Genetics and Lineberger Comprehensive Cancer Center, University of North Carolina, Chapel Hill, NC; Department of Ecology and Evolutionary Biology, Cornell University, Ithaca, NY; Island Conservation, Santa Cruz, CA; Re:wild, Puerto Ayora, Galápagos Islands, Ecuador; Institute for Genome Sciences and Society, Texas A&M University, College Station, TX; Institut des Sciences de l’Évolution Montpellier, Université de Montpellier, Montpellier, France

**Author notes:** Corresponding authors: Andrew P Morgan, Fernando Pardo-Manuel de Villena, 5049C Genetic Medicine Building, Department of Genetics, University of North Carolina, Chapel Hill NC 27599-7264. These authors contributed equally (Andrew P Morgan, Jonathan J Hughes). Department of Medicine, Duke University Hospital, Durham, NC. Independent scientist (John P Didion). Co-senior authors (Jeremy B Searle, Fernando Pardo-Manuel de Villena).

## Abstract

House mice (*Mus musculus*) have spread globally as a result of their commensal relationship with humans. In the form of laboratory strains, both inbred and outbred, they are also among the most widely-used model organisms in biomedical research. Although the general outlines of house mouse dispersal and population structure are well known, details have been obscured by either limited sample size or small numbers of markers. Here we examine ancestry, population structure and inbreeding using SNP microarray genotypes in a cohort of 814 wild mice spanning five continents and all major subspecies of *Mus*, with a focus on *M. m. domesticus*. We find that the major axis of genetic variation in *M. m. domesticus* is a south-to-north gradient within Europe and the Mediterranean. The dominant ancestry component in North America, Australia, New Zealand and various small offshore islands is of northern European origin. Next we show that inbreeding is surprisingly pervasive and highly variable, even between nearby populations. By inspecting the length distribution of homozygous segments in individual genomes, we find that inbreeding in commensal populations is mostly due to consanguinity. Our results offer new insight into the natural history of an important model organism for medicine and evolutionary biology.

## Introduction

The house mouse (*Mus musculus*) is among the best established model organisms for genetic studies in mammals, having been utilized in biomedical research for decades and leading to many fundamental discoveries (Vandenbergh 2000; Guénet and Bonhomme 2003). Furthermore, mice have been used successfully to explore broad evolutionary questions of speciation and hybrid zones, adaptation, and karyotype evolution amongst others (Sage *et al*. 1993; Phifer-Rixey and Nachman 2015). Their status as a laboratory model has helped develop them as a staple of evolutionary biology research, given the ease with which wild mice can be reared in captivity, as well as the now abundant genomic resources available for the study of mice.

The earliest recognized instances of *Mus musculus* have been traced back approximately 1 mya to central Asia and Northern India (Boursot *et al*. 1993; Suzuki and Aplin 2012). *Mus musculus* then began to diverge 0.25– 0.5 mya (Bonhomme and Searle 2012; Phifer-Rixey *et al*. 2020) into three major subspecies, and with human-mediated movements have gained large ranges: *M. m. domesticus* (Western Europe, the Americas, Africa and Australasia), *M. m. musculus* (Eastern Europe and Northern Asia), and *M. m. castaneus* (Southern Asia). These subspecies can be distinguished morphologically (Macholán 1996), with biochemical and cytogenetic markers (Bonhomme *et al*. 1984), with mitochondrial DNA (mtDNA) (Prager *et al*. 1998) and now on the basis of genome-wide SNP or sequence data (Yang *et al*. 2009). Several other subdivisions of *Mus musculus* have been proposed on the basis of morphology, karyotype, mitochondrial haplotypes or microsatellite data (Sage 1981; Prager *et al*. 1998; Piálek *et al*. 2005; Suzuki *et al*. 2013; Hardouin *et al*. 2015). *M. m. molossinus* is a hybrid of *M. m. musculus* and *M. m. castaneus* endemic to Japan (Yonekawa *et al*. 1988). A mitochondrial lineage in the Arabian Peninsula and Madagascar has been labelled *M. m. gentilulus* (Prager *et al*. 1998; Duplantier *et al*. 2002). The taxonomic status of a group of populations in Iran, Pakistan, Afghanistan and northern India – overlapping with the ancestral range of house mice – is somewhat uncertain. Some were historically classified as *M. m. bactrianus* (Schwarz and Schwarz 1943) *but they are now thought to represent deep divisions within M. m. castaneus* (Rajabi-Maham *et al*. 2012; Hamid *et al*. 2017). In this manuscript we refer to this group as *M. musculus* undefined, and include possible *M. m. gentilulus* specimens in this category.

The commensal relationship of the house mouse with humans, a characteristic which is absent in their closest relatives *M. spretus, M. spicilegus*, and *M. macedonicus*, has facilitated their global dispersal (Chevret *et al*. 2005; Weissbrod *et al*. 2017). The earliest fossil evidence suggests the commensalism of mice with humans began around 15 kya in the Levant, followed by expansion into northern and western Europe as a result of increased human trade and settlement (Cucchi *et al*. 2020). Mus musculus experienced far reaching range expansion during the age of European exploration and colonization, allowing them to establish populations on oceanic islands and in the New World alongside human colonists (Boursot *et al*. 1993; Gabriel *et al*. 2010; Hardouin *et al*. 2010; Bonhomme and Searle 2012).

One of the most interesting and unusual features of European *M. m. domesticus* is the number and frequency of chromosomal rearrangements that segregate in natural populations. Numerous Robertsonian metacentric chromosomes (centromere-to-centromere fusions) have been described (Capanna *et al*. 1976; Piálek *et al*. 2005). The processes that generate these rearrangements and allow them to persist in populations, despite adverse effects on reproduction in heterozygotes, have been the subject of intense study over the past 50 years (Garagna *et al*. 2014; Giménez *et al*. 2017). Populations with a characteristic complement of metacentric chromosomes are referred to as “chromosomal races.” In this paper, we include the data for the chromosomal races as part of our broader study of house mouse. A companion paper (Hughes *et al*., in preparation) will focus specifically on the genomics of chromosomal variation in the house mouse.

Given the diversity of habitat, evolutionary history, and phenotype of wild house mouse populations, as well as their close relationships to humans, they are a wellspring for potential scientific inquiry. However, the majority of studies focus on laboratory mice that stem from deliberate inbreeding (Phifer-Rixey and Nachman 2015). It is difficult to capture the true scale of genetic and phenotypic diversity of wild populations with what constitutes a small global sampling (Keane *et al*. 2011). The genomes of so-called “classical laboratory strains” of mice are a mosaic of the three major subspecies, with on average 92% *M. m. domesticus* ancestry (Yang *et al*. 2007, 2011; Didion and de Villena 2013). A smaller collection of inbred strains are derived from wild-caught ancestors from each of the major subspecies. A more complete understanding of the house mouse system necessarily requires a greater wealth of genetic information from wild populations than is presently available.

Here we make use of genome-wide SNP genotypes obtained with two microarray platforms to characterize the ancestry and population structure of a large survey of wild-caught mice from around the globe. We characterize the utility of these arrays for population-genetic analysis by comparison to whole-genome sequence data from a representative subsample of mice. We explore the relationship between population structure, inbreeding and characteristics of where mice are living. Our study represents one of the largest and most comprehensive descriptions of the genetic structure of house mouse populations to date.

## Materials and methods

### Sample collection

Mice were collected at 268 locations in 33 countries (recorded in **Table S1**) between 1990 and 2015. All trapping and euthanasia was conducted according to the animal use guidelines of the institutions with which providers were affiliated at the time of collection. Trapping on Southeast Farallon Island was carried out during two seasons, in 2011 and 2012; and on Floreana during a single season in 2012. Particular sampling effort was dedicated to regions with high karyotypic diversity such as Greece, the Swiss-Italian Alps and the island of Madeira, where hybridization between chromosomal races is known to occur (Britton-Davidian *et al*. 2000; Hauffe *et al*. 2012).

Samples of one or more tissues (including tail, liver, spleen, muscle and brain) from each individual were shipped to the University of North Carolina during an eight-year period (2010-2017). We assigned each individual a unique identifier as follows: `CC:LLL_SSSS:DD_UUUU` (`C` = ISO 3166-1-alpha2 country code; `L` = locality designation; `S` = nominal subspecies; `D` = diploid chromosome number and `U` = sequential numeric ID).

Genotypes for some individuals in this study have been used in prior studies on a selfish genetic element in European *M. m. domesticus* (Didion *et al*. 2016) and/or on an introduced population of house mice in New Zealand (predominantly *M. m. domesticus* but with genomic contributions from *M. m. musculus* and *M. m. castaneus*) (Veale *et al*. 2018).

### DNA preparation and genotyping

Whole-genomic DNA was isolated from tissue samples using Qiagen Gentra Puregene or DNeasy Blood & Tissue kits according to the manufacturer’s instructions. Genotyping was performed using either the Mega Mouse Universal Genotyping Array (MegaMUGA) or its successor, GigaMUGA (GeneSeek, Lincoln, NE). Genotypes were called using Illumina BeadStudio (Illumina Inc, Carlsbad, CA). Only samples with *<* 10% missing calls were retained for analysis.

Sample sexes were confirmed by comparing the number of non-missing calls on the Y chromosome to the number of heterozygous calls on the X chromosome (**Figure S1**).

### Whole-genome sequencing

Whole-genome sequence (WGS) data was obtained from the European Nucleotide Archive for 136 mice representing natural populations of *M. m. domesticus* (Europe: PRJEB9450 (Harr *et al*. 2016); North America: PRJNA397406 (Phifer-Rixey *et al*. 2018); Gough Island: PRJNA587779 (Wang *et al*. 2017), PRJNA352398 (Gray *et al*. 2015)), *M. m. castaneus* (PRJEB2176) (Halligan *et al*. 2013), M. m. musculus (PRJEB14167, PRJEB11742), *M. spretus* (PRJEB11742) (Harr *et al*. 2016) plus one *M. spicilegus* (PRJEB11513) (Neme and Tautz 2016) individual.

New sequencing data was generated for 5 individuals (EC:FLO_STND:40_6255, EC:FLO_STND:40_3043, US:SEF_STND:40_3023, US:SEF_STND:40_3023, UK:INA_STND:40_278). Sequencing was performed at the University of North Carolina High Throughput Sequencing Facility. Whole genomic DNAs were sheared by ultrasonication and the resulting fragments were size selected to target size 350 bp using a PippinPrep system (Sage Sciences, Beverly, MA).The UNC High Throughput Sequencing Facility generated sequencing libraries using Kapa DNA Library Preparation Kits (Kapa Biosystems, Wilmington, MA). Each library was run on its own lane of a HiSeq4000 (Illumina Inc, Carlsbad, CA), and generated 150 bp paired-end reads.

### Genotype calling from whole-genome sequencing

Raw reads were aligned to the mm10 mouse reference genome with `bwa mem` v0.7.15; optical duplicates were marked with `samblaster` v0.1.22 and excluded from further analyses. Genotypes were called at 64 886 target sites corresponding to known SNPs with unique probe sequences on both MegaMUGA and GigaMUGA arrays (see below) using `GATK` v4.1.0.0 Handsaker *et al*. (2011). Joint genotype calling was performed across all samples using the `HaplotypeCaller` - `CombineGVCFs` - `GenotypeGVCFs` workflow. Full details can be found in the scripts on Github at https://github.com/andrewparkermorgan/wild_mouse_genetic_survey.

### Genotype merging and filtering

Genotypes from the two array platforms and WGS were merged into a single matrix as follows. Karl Broman’s re-annotation of the MUGA family of arrays (https://github.com/kbroman/MUGAarrays) was used as a starting point. For the 64 886 sites targeted on both MegaMUGA and GigaMUGA, a single representative probe on each array was selected (some sites are targeted by multiple probes). Alleles were swapped such that all genotypes were reported on the positive strand and prepared for merging.

Genotypes were successfully called in WGS at 62 489 (96.3%) of the target sites. Of these, 57 945 (89.3%) were biallelic among WGS samples and had the same alternate allele as targeted on the arrays. These sites represent the intersection of the two array platforms and WGS. Array and WGS genotypes were then merged into a single VCF file with `bcftools isec` v1.9.

The merged genotype matrix has dimensions 57 945 sites *×* 814 individuals with final genotyping rate 97.1%. It is available on Dryad at https://doi.org/10.5061/dryad.ncjsxkswt.

### Relatedness

Cryptic relatives were identified on the basis of autosomal genotypes with `akt kin` v.3beb346 (Arthur *et al*. 2017). Hereafter we denote the *n × n* kinship matrix **K** and its entries *k*_*ij*_ where *i, j* are indices for individuals and *i* ≠ *j*. Kinship estimation was performed separately in each taxon (*M. m. domesticus, M. m. musculus, M. m. castaneus, M. musculus* undefined, *M. spretus*) because allele frequencies are highly stratified between these groups. For each pair with kinship coefficient *>* 0.10, one member was removed at random to yield a set of 651 putatively-unrelated individuals. We confirmed that qualitative results from analyses of population structure by both PCA and `ADMIXTURE` (see below) were insensitive to the exact composition of the putatively-unrelated subset by re-running analyses on 10 random unrelated subsets. Results from this exercise are shown in **Figure S2** and **Figure S3**.

### Analyses of population structure

Principal components analysis (PCA) was performed with `akt pca` v.3beb346 (Arthur *et al*. 2017), using only autosomal sites with genotyping rate *>* 90% and minor-allele frequency (MAF) *>* 1%. PCs were first estimated from a subset of individuals selected as follows to maximize geographic representation while balancing sample size across geographic locations. Starting from the list of 651 putatively-unrelated individuals, we randomly selected up to 10 from each geographic location. This yielded 255 *M. m. domesticus*, 24 *M. m. musculus*, 39 *M. m. castaneus*, 25 *M. musculus* undefined and 6 *M. spretus* individuals. Genotypes from the entire cohort were then projected onto the PCs estimated in the representative subset.

A Y chromosome phylogenetic tree was inferred from 19 Y-linked sites with *<* 10% missing genotypes (among males only), using neighbor-joining as implemented in the `bionjs()` function in the `ape` package v5.1 (Paradis *et al*. 2004). The tree was re-rooted using *M. spicilegus* as outgroup.

Further ancestry analyses within *M. m. domesticus* were performed by running `ADMIXTURE` in unsupervised mode with *K* = 2, …, 5 components on a genotype matrix including only the 56 201 autosomal sites. As for PCA, we estimated component-specific allele frequencies (the *P* matrix) using a representative subset of individuals and estimated ancestry proportions (the *Q* matrix) with the values of *P* held fixed, to avoid distortion due to over-sampling of certain locations. `ADMIXTURE` is best understood as a “grade-of-membership” model (Erosheva 2006) that models allele frequencies in each individual as a mixture of allele frequencies in *K* putative source populations. For this interpretation to hold, the true source populations must be well differentiated and present in the dataset at hand. Furthermore, other demographic processes with strong effects on allele frequencies, such as population bottlenecks or rapid expansion, must not have occurred. These assumptions are often violated in practice, including in our study. Under these conditions the notion of an “optimal” value of *K* loses its meaning (Lawson *et al*. 2018). We treat `ADMIXTURE` as a descriptive tool that reveals different levels of population structure at increasing values of *K* and do not attempt to find an optimal value for *K*.

A “population tree” was produced with `TreeMix` v1.12r231 (Pickrell and Pritchard 2012). *M. spretus* individuals from Spain were used as outgroup to root trees. Runs were performed with *m* = 0, …, 3 gene-flow edges. Outgroup *f*_3_-statistics were calculated with the `threepop` utility and four-population *f*_4_-statistics for admixture graphs with the `fourpop` utility, both included in the `TreeMix` suite; standard errors were calculated by block jackknife over blocks of 500 sites.

### Inbreeding

Individual inbreeding coefficients 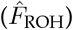 were estimated as the fraction of the autosomes covered by runs of homozygosity (ROH). This has previously been shown to be a reasonably consistent estimator of recent inbreeding (McQuillan *et al*. 2008). ROH were identified separately in each individual using `bcftools roh` v1.9 (Narasimhan *et al*. 2016) with taxon-specific allele frequencies (estimated over unrelated individuals only in each of *M. m. domesticus, M. m. musculus, M. m. castaneus*), constant recombination rate 0.5 cM/Mb (Liu *et al*. 2014), and arbitrarily assigning array genotypes a Phred-scale GT = 30 (corresponding to error rate 0.001) per the authors’ recommendations. Other HMM parameters were left at their defaults after experimenting with a wide range of parameter settings, and finding the root mean squared error (relative to estimates of 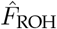 from WGS data) to be insensitive to these parameters. We then excluded segments *<* 2 cM in length. Mean and median length of remaining putative ROH segments were 13.2 Mb (287 sites) and 6.8 Mb (151 sites) respectively. Between-group differences in inbreeding coefficients were evaluated using linear models with logit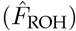 as the response variable. For these analyses we excluded 4 mice (SH:GOU_STND:40_6256, SH:GOU_STND:40_6257, SH:GOU_STND:40_6258, SH:GOU_STND:40_6259) that were members of a laboratory colony established from wild-caught founders from Gough Island, but had been partially inbred in the laboratory.

### Maps and plotting

Figures were created with the `R` packages `ggplot2` v3.3.1 (Wickham 2016), `ggbeeswarm` v0.7.0 (https://github.com/eclarke/ggbeeswarm), `viridis` v0.5.1 (https://CRAN.R-project.org/package=viridis), `maps` v3.3.0 (https://CRAN.R-project.org/package=maps), `cowplot` v0.9.3 (https://CRAN.R-project.org/package=cowplot), `popcorn` v0.0.2 (https://github.com/andrewparkermorgan/popcorn) and `mouser` v0.0.1 (https://github.com/andrewparkermorgan/mouser).

## Results

Our dataset consists of 814 mice (**Figure 1, Table S1**) spanning the three principal subspecies of the house mouse found across its global range (*M. m. domesticus, M. m. musculus, M. m. castaneus* plus 2 early-generation hybrids), the outgroup species *M. spretus* and *M. spicilegus*, and a group with ill-defined ancestry provisionally labelled “*M. musculus* undefined.” Our collection includes 218 mice carrying Robertsonian translocations, representing 37 chromosomal races with diploid number (2*N*) between 22 and 39. The majority of specimens were genotyped with Illumina Infinium SNP arrays (Morgan *et al*. 2016) and the remainder were obtained from published whole-genome sequencing (WGS) data (Pezer *et al*. 2015; Harr *et al*. 2016; Neme and Tautz 2016). Sample sizes by taxon and genotyping platform are summarized in **Table 1**. Briefly, the two array datasets were merged on overlapping markers, and WGS samples were added by calling genotypes at array sites. WGS data were also used to identify and remove problematic sites. Full details of data processing and quality control are provided in the **Materials and methods**. The final genotype matrix comprises 57 945 sites spanning 2390 Mb (99.4%) and 164 Mb (97.9%) of autosomal and X chromosome sequence, respectively; corresponding to 1296 cM (89.9%) and 70.6 cM (88.9%) on the genetic map, respectively. Median spacing between adjacent sites is 28.9 kb (median absolute deviation [MAD] 25.9 kb) on the autosomes and 51.1 kb (MAD 51.8 kb) on the X chromosome (**Figure S4**). The mutation spectrum is biased towards transitions (Ti:Tv = 4.71) as a result of technical constraints imposed by the array platform, as described elsewhere (Morgan *et al*. 2016).

**Table 1:** Count of samples used in this study by taxon and genotyping platform.

|  | GigaMUGA | MegaMUGA | WGS | Total |
| --- | --- | --- | --- | --- |
| *M. m. domesticus* | 143 | 468 | 99 | 710 |
| *M. m. musculus* | 0 | 6 | 22 | 28 |
| *M. m. castaneus* | 0 | 29 | 10 | 39 |
| *M. musculus* undefined | 0 | 27 | 0 | 27 |
| *M. spretus* | 0 | 0 | 7 | 7 |
| *M. spicilegus* | 0 | 0 | 1 | 1 |
| *dom-mus* hybrid | 0 | 2 | 0 | 2 |
| **Total** | 143 | 532 | 139 | 814 |

**Figure 1:**
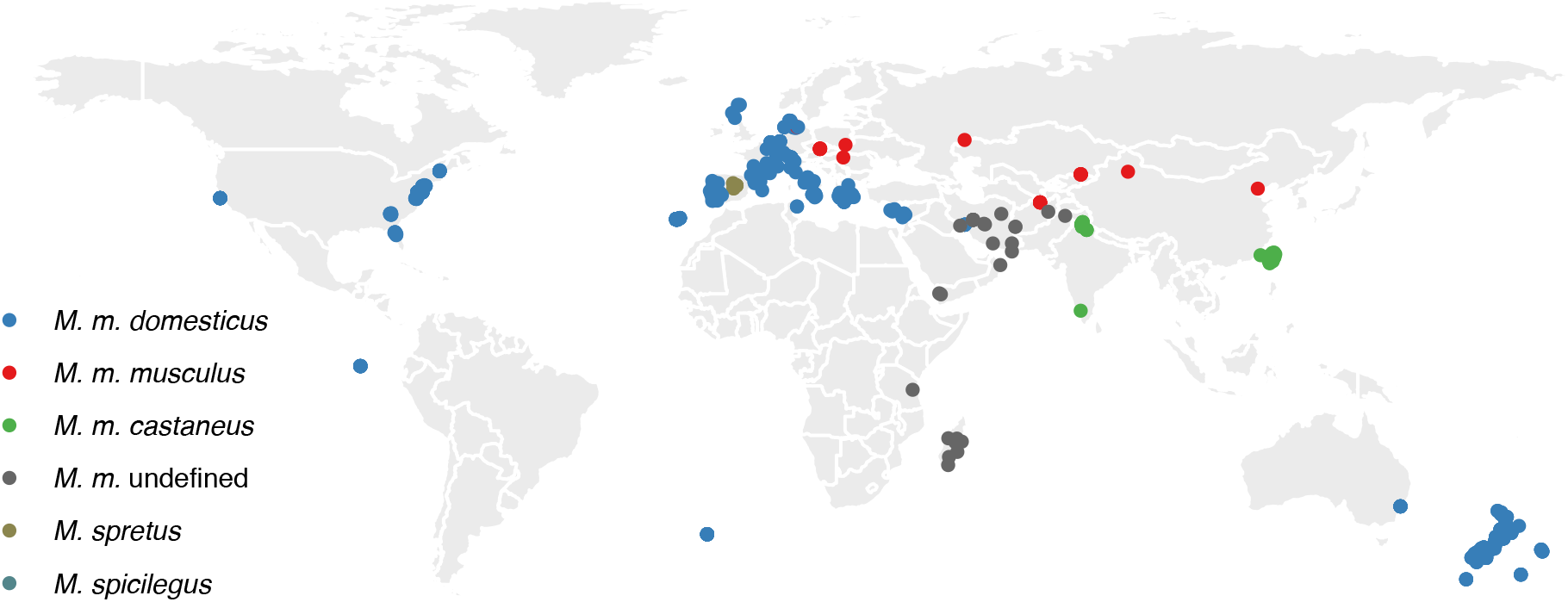
Geographic distribution of wild house mice used in this study. One point per individual, coloured by species or subspecies of origin.

The genotyping arrays used in this study have a number of design properties that constrain their use for population genetics studies (Morgan *et al*. 2016). The number of markers that are polymorphic within *M. m. domesticus* (54 929, 94.8%) is much greater than within *M. m. musculus* (24 330, 42.0%) or *M. m. castaneus* (32 633, 56.3%) (**Table 2**). The site frequency spectrum, expected to have an approximately exponential shape under a wide range of demographic scenarios (Fu 1995), is strongly biased towards intermediate frequencies (**Figure S5**), especially within *M. m. domesticus*. As a consequence, many summary statistics that can be derived from the site frequency spectrum (pairwise diversity, Watterson’s *θ*, Tajima’s *D*) do not have interpretable values. The effects of SNP ascertainment bias on population genetic studies have been explored in detail elsewhere (for example Lachance and Tishkoff 2013). It suffices to say here that array genotypes are still useful for analysing population structure and ancestry if used with appropriate caution.

**Table 2:** Number and proportion of markers polymorphic in each taxon, by chromosome type (A = autosomal, X = X-linked, Y = Y-linked, M = mitochondrial).

|  | **A**<br>(56201) | **X**<br>(1711) | **Y**<br>(19) | **M**<br>(14) | **Total**<br>(57945) |
| --- | --- | --- | --- | --- | --- |
| *M. m. domesticus* | 53464 (95.1%) | 1434 (83.8%) | 17 (89.5%) | 14 (100%) | 54929 (94.8%) |
| *M. m. musculus* | 23957 (42.6%) | 360 (21%) | 3 (15.8%) | 10 (71.4%) | 24330 (42%) |
| *M. m. castaneus* | 32077 (57.1%) | 539 (31.5%) | 8 (42.1%) | 9 (64.3%) | 32633 (56.3%) |
| *M. musculus* undefined | 38270 (68.1%) | 868 (50.7%) | 7 (36.8%) | 8 (57.1%) | 39153 (67.6%) |
| *M. spretus* | 2027 (3.6%) | 23 (1.3%) | 0 (0%) | 7 (50%) | 2057 (3.5%) |

### Microarray genotypes recover subspecies relationships

To confirm nominal subspecies assignments for our samples, we performed principal components analysis (PCA) on autosomal genotypes for a representative subset of 374 individuals. Three major clusters are clearly observed, corresponding to the three principal subspecies (**Figure 2A**). Both *M. m. domesticus* and *M. m. musculus* form relatively tight clusters in the top PCs, while *M. m. castaneus* forms two sub-clusters corresponding to mice from Taiwan and India, respectively. This is consistent with existing evidence for greater variation within *M. m. castaneus* compared to the other two major subspecies (eg. Geraldes *et al*. 2008). *M. musculus* undefined specimens project near the *M. m. castaneus* cluster, consistent with prior reports. To directly examine phylogenetic relationships, we chose the only non-recombining locus for which our genotype matrix contains sufficient data – the Y chromosome. A neighbor-joining tree from Y-linked sites recovers the split between *M. m. domesticus* and all other groups, as well as at least one instance of intersubspecific Y chromosome introgression from a non-*M. m. domesticus* into a *M. m. domesticus* individual from Porto Santo (**Figure 2B**). *M. m. castaneus* and *M. m. musculus* cannot be distinguished in this analysis due to a lack of subspecies-diagnostic markers on the Y chromosome in the merged genotype matrix.

**Figure 2:**
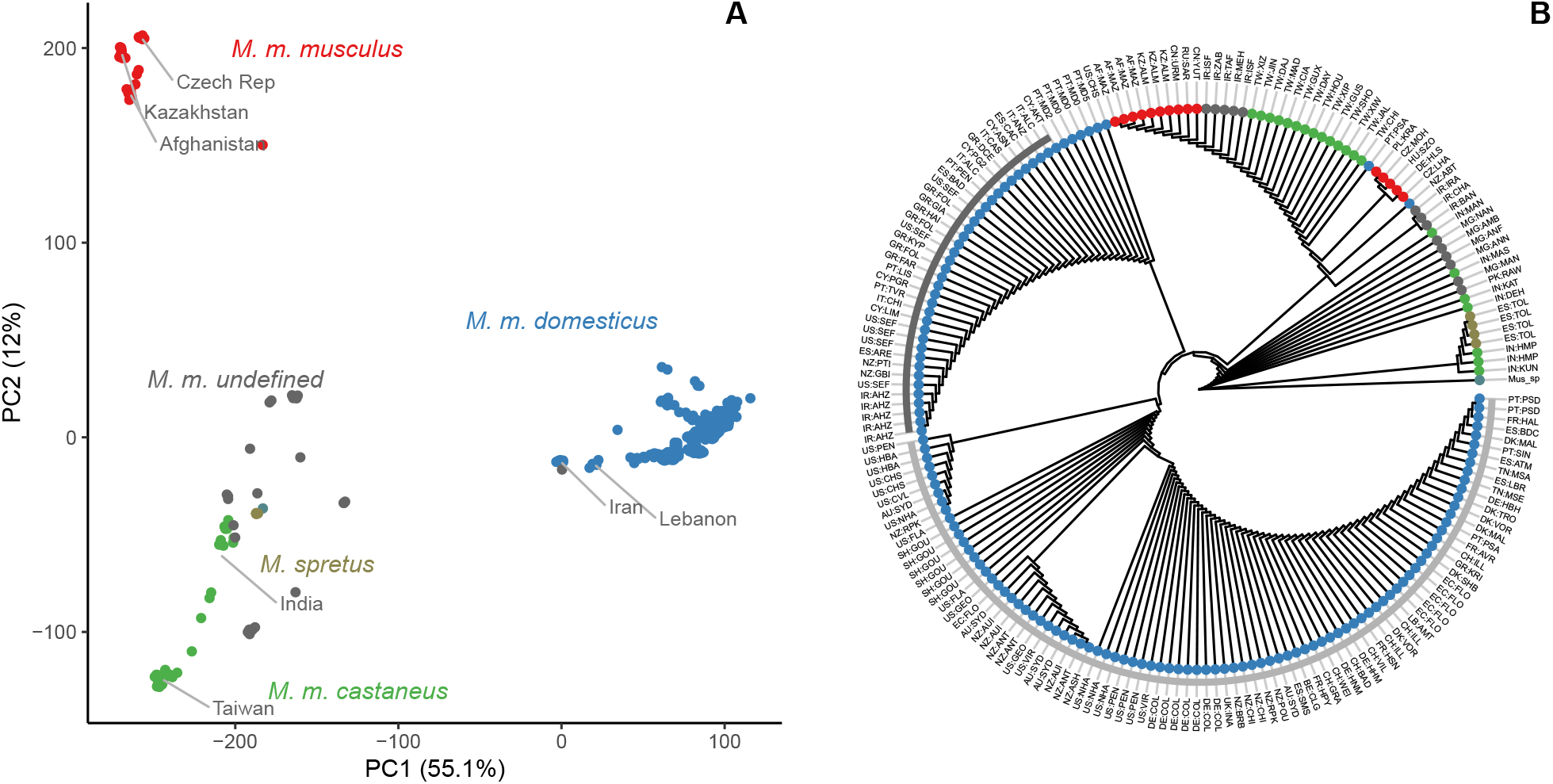
Array genotypes discriminate between *Mus musculus* subspecies. (**A**) PCA on autosomal genotypes. (**B**) Neighbor-joining tree of Y chromosome haplotypes from representative individuals. Two distinct haplogroups within *M. m. domesticus* are marked with dark and light grey bars, respectively.

### Population structure in European *M. m. domesticus* follows a latitudinal gradient

The primary focus of our work is population structure within *M. m. domesticus*. We again performed PCA anchored by a representative subset of individuals. Several patterns are evident. First, the major axis of variation is a south-to-north gradient across Europe and western Eurasia. We emphasize this by assigning individuals to three major geographic groups in **Figure 3A**. Plotting the first PC against location makes the spatial gradient clear (**Figure 3B**). Second, individuals cluster by location within broader geographic groups (**Figure 3C-E**). Third, *M. m. domesticus* that have dispersed outside Eurasia tend to cluster with northern European populations.

**Figure 3:**
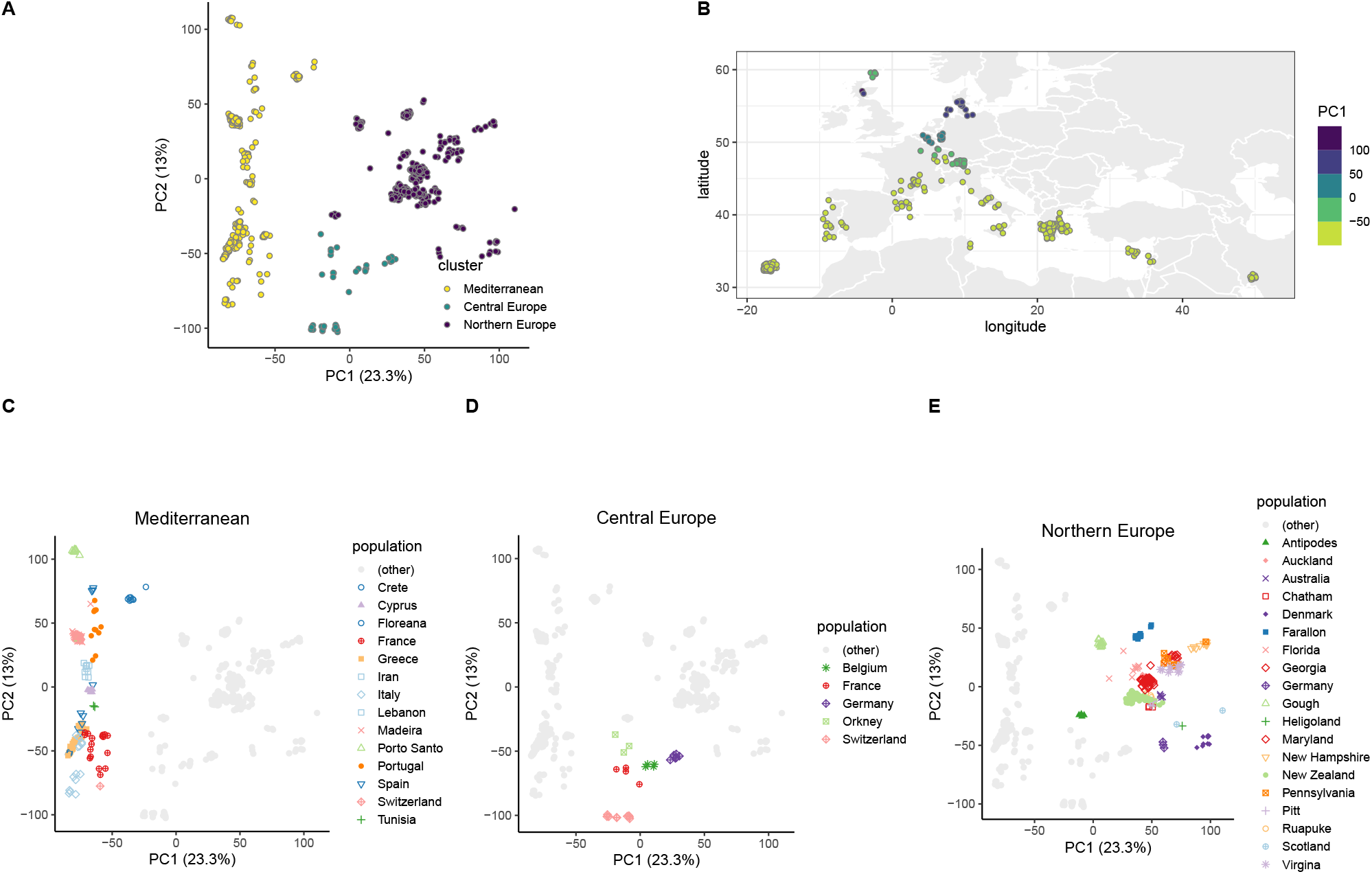
Population structure in *M. m. domesticus*. (**A**) PCA on autosomal genotypes (PC1 *vs* PC2 shown), with individuals coloured according to membership in broad geographic groups named after predominant European areas represented in each. (**B**) PC1 coordinate versus sampling location. (**C-E**) Same PCA plot as in panel A, but emphasizing sub-structure within large-scale geographic groups. Points coloured according to population of origin.

We also used `ADMIXTURE` (Alexander *et al*. 2009) to estimate ancestry proportions with increasing numbers of ancestry components (**Figure S6**). This analysis shows that ancestry profiles within geographically-defined populations are generally cohesive. At the most coarse scale (*K* = 2), we again see a broad division between populations from the Mediterranean and the Iberian Peninsula (light blue component) and those from northern Europe and the New World (dark blue component) (**Figure 4**). There is a similar result to the `TreeMix` analysis (**Figure 5A**) with the two groupings again evident. The southern European grouping represents well the known initial colonization of Europe by *M. m. domesticus* based on archaeological data (Cucchi *et al*. 2005) (**Figure 5B**). The colonization route and derivation of the northern and central European groupings is less clear.

**Figure 4:**
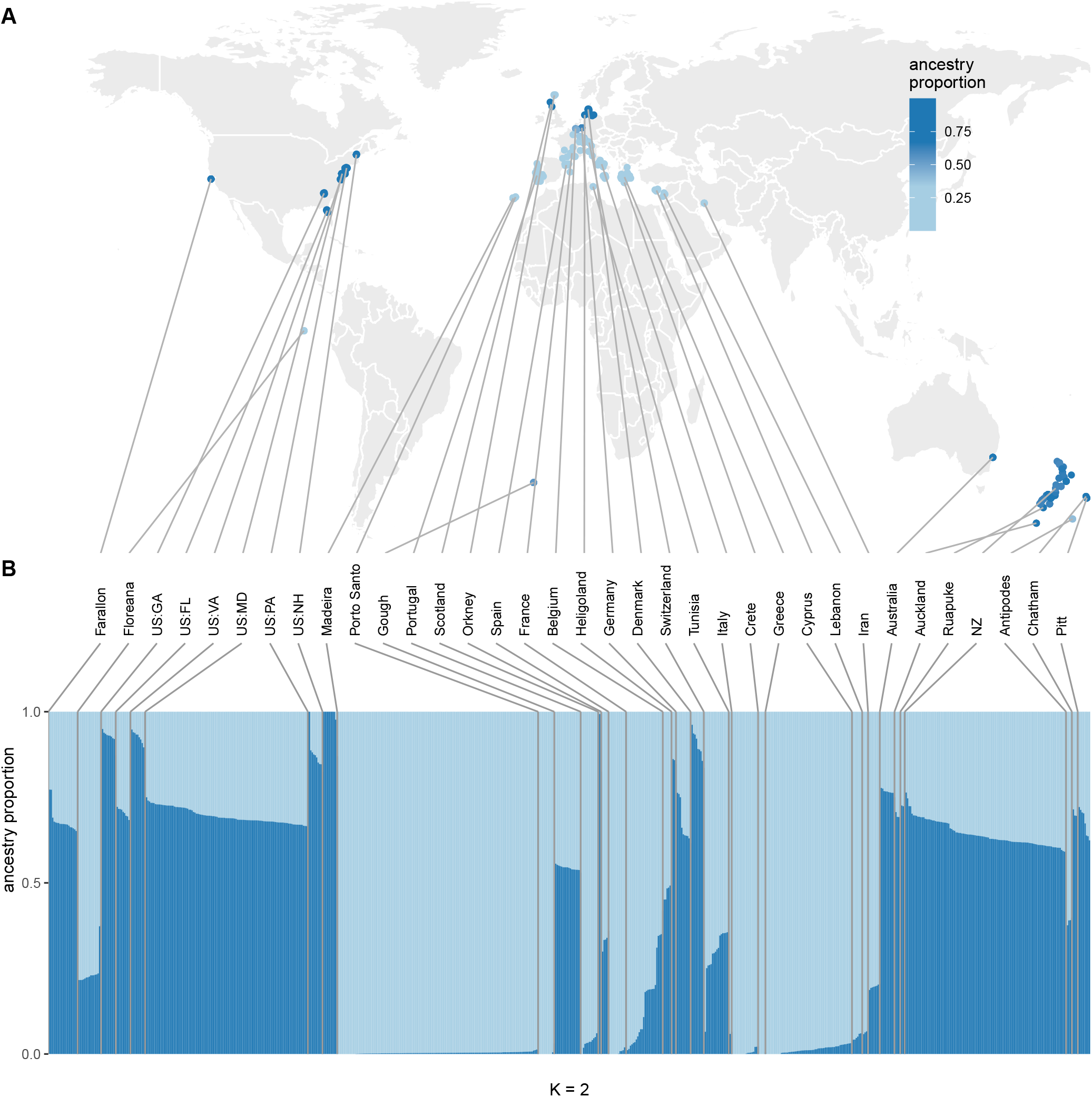
Ancestry decomposition with `ADMIXTURE`. (**A**) Individuals plotted by location, coloured according to ancestry proportion. (**B**) Ancestry proportions estimated with `ADMIXTURE` for *K* = 2. Population labels shortened for legibility: US:GA = Georgia, US:FL = Florida, US:VA = Virginia, US:MD = Maryland, US:PA = Pennsylvania, US:NH = New Hampshire, NZ = New Zealand.

**Figure 5:**
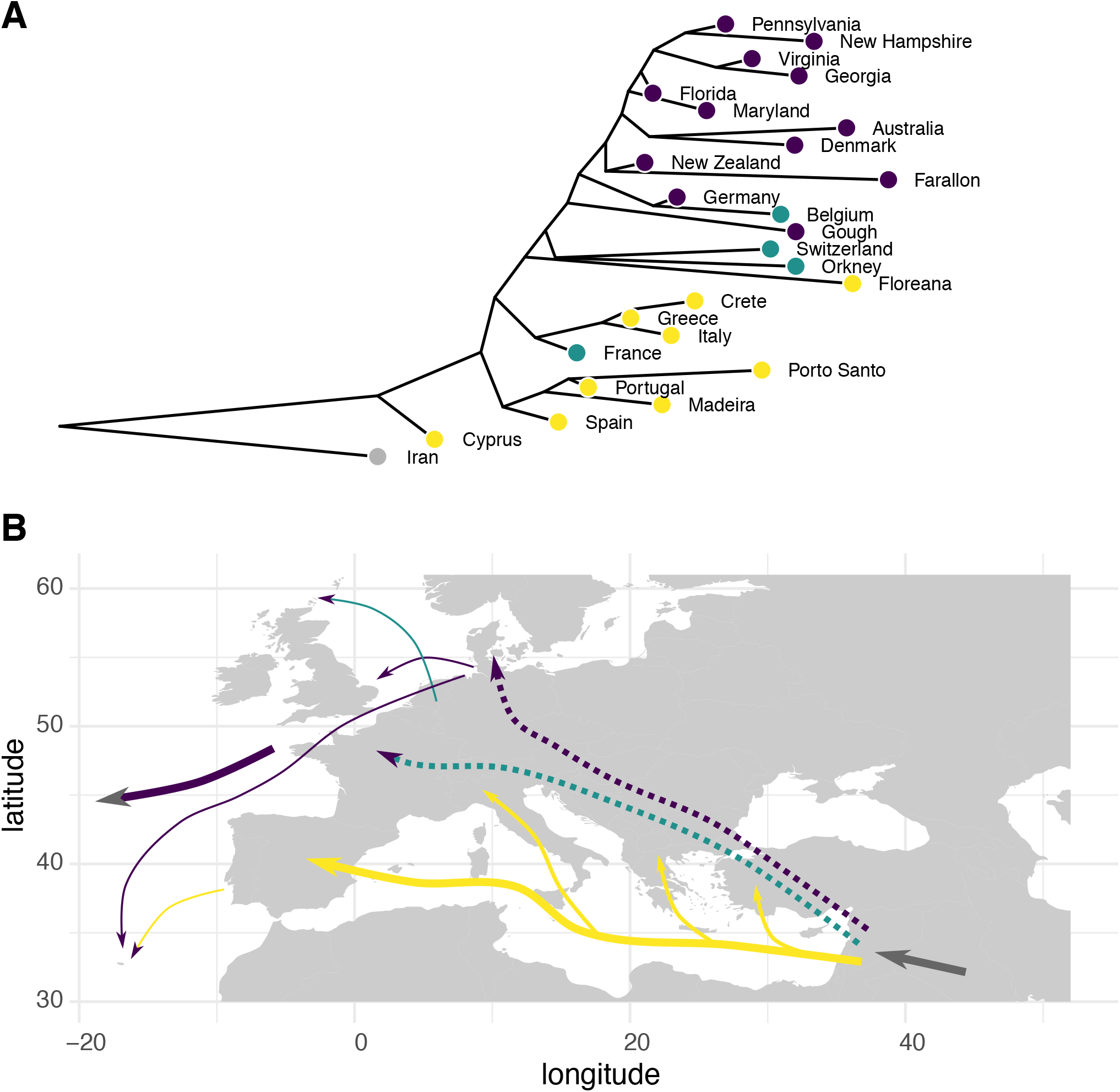
Origins of European *M. m. domesticus* populations. (**A**) Population tree inferred by `TreeMix`, with populations coloured by broad geographic group as in **Figure 3**. (**B**) Model for dispersal of *M. m. domesticus* across Europe, based on current distributions of the different groups and archaeological interpretations (Cucchi *et al*. 2005). The derivation and route of colonization of the northern and central European groupings (shown in green and purple) is unclear (as indicated by dashed lines).

### Inbreeding is pervasive but highly variable

The deme structure of natural mouse populations predisposes to inbreeding, and several mechanisms of kin recognition and inbreeding avoidance based on scent have also been proposed (Yamazaki *et al*. 1976; Hurst *et al*. 2001; Sherborne *et al*. 2007). Previous estimates of inbreeding in wild mice vary widely within and between populations, with average between 0.2 − 0.5 based on microsatellites (Ihle *et al*. 2006; Hardouin *et al*. 2015) or SNPs (Laurie *et al*. 2007). We used the proportion of the autosomes contained in runs of homozygosity (ROH) as an estimator of the inbreeding coefficient 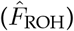, as it has been shown to be relatively powerful and less prone to bias due to stratification of allele frequencies than other estimators (Keller *et al*. 2011). 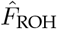 values vary widely within and between subspecies, shown in **Figure 6A** (ANOVA: *F*_2,760_ = 4.88, *p* = 7.8 *×* 10^−3^). The highest values are observed in *M. m. domesticus* and the lowest in *M. m. castaneus* (*t* = 2.93, *p*_adj_ = 9.8 *×* 10^−3^ by Tukey’s post-hoc method). Among females, who carry two copies of the X chromosome, homozygosity on the X chromosome is positively correlated with 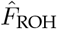 estimated from autosomal sites (**Figure 6B**) as expected. Inbreeding estimates tend to be consistent within geographically-defined groups, and there is significant heterogeneity across these groups (ANOVA: *F*_24,676_ = 14.3, *p <* 10^−6^) (**Figure 6C**). Some of the highest 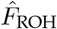 values are on some of the smallest islands (Heligoland, Orkney, Farallon, Floreana), but there are also small islands with much lower values (Madeira, Porto Santo) (**Figure 6C**). Although there is a nominal difference in mean 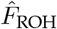 between mice with standard versus Robertsonian karyotypes (*t* = 14.1, *p* = 1.8 *×*10^−4^), the distribution of 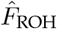 among mice with standard karyotypes clearly spans that of Robertsonian mice (**Figure 6D**). The relationship between karyotype variation and inbreeding is explored further in the companion manuscript (Hughes *et al*., in preparation).

**Figure 6:**
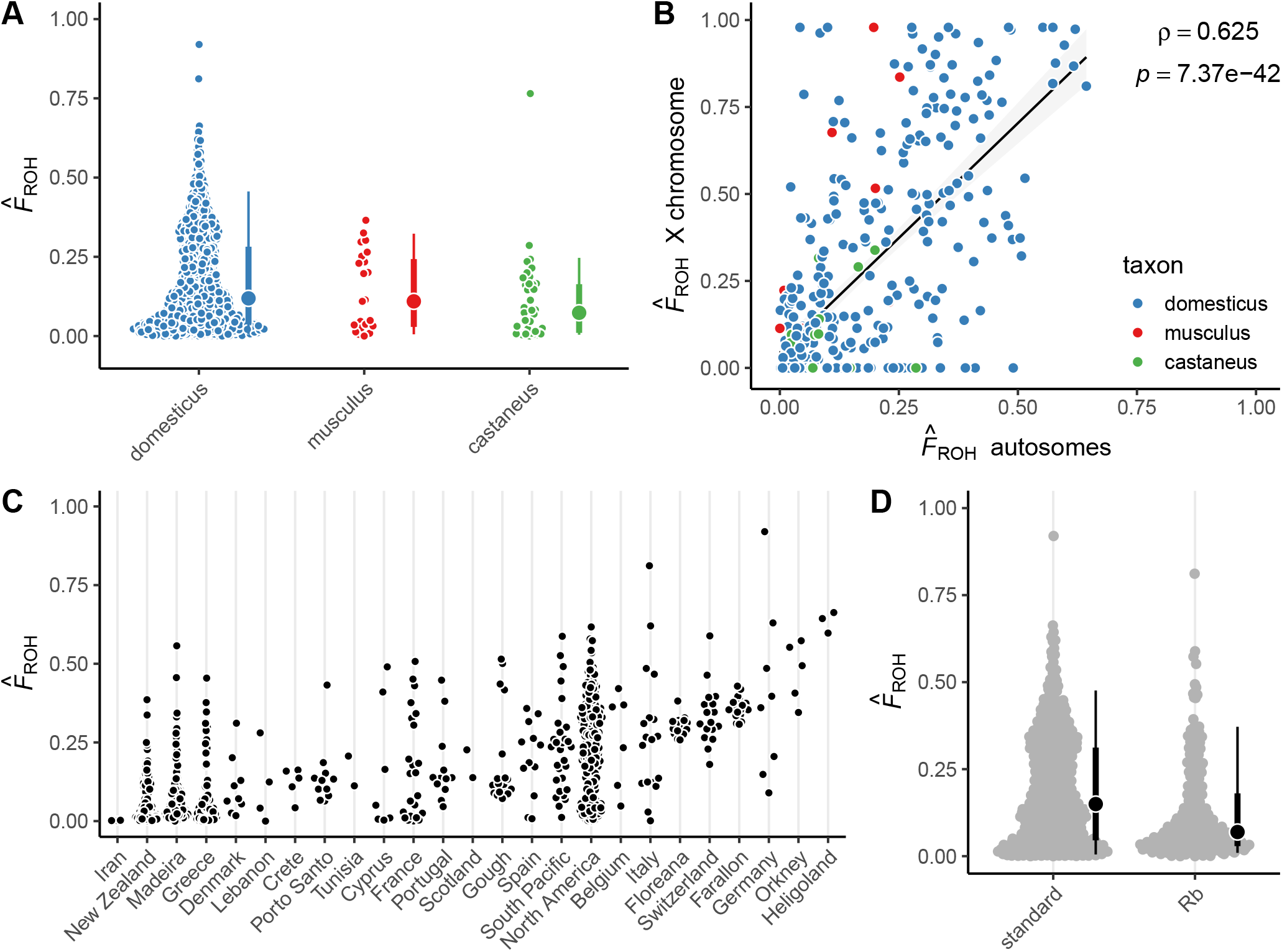
Inbreeding across mouse subspecies and populations. (**A**) Inbreeding coefficients estimated from autosomal genotypes, by nominal subspecies of origin. Open dot, median; thick bar, interquartile range; thin bar, 5^th^ - 95^th^ percentiles. (**B**) Autosomal *vs* X chromosome inbreeding coefficients for 374 female mice. Black line, linear regression through all points; grey band, 95% confidence region. Spearman’s (rank) correlation and corresponding *p*-value shown in upper right. (**C**) Inbreeding coefficients by population within *M. m. domesticus*. Populations are sorted by mean 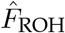. (**D**) Comparison of inbreeding coefficients in *M. m. domesticus* with standard *vs* Robert-sonian (Rb) karyotypes.

Runs of homozygosity reflect sharing of chromosomal segments identical by descent between an individual’s parents (ignoring rare instances of uniparental disomy). When the common ancestor from which these segments were inherited is deep in the past, the expected length of the shared segment is shorter; when the common ancestor is more recent, the shared segment is on average longer. Deeper pedigree connections may arise from small historical population sizes or founder effects. More recent pedigree connections reflect mating between close relatives; that is, consanguinity. A genome-wide estimator of homozygosity such as 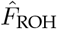 thus comprises a mixture of shared segments from pedigree connections of varying depth. The distribution of ROH segment lengths is informative for both recent and historical demographic processes.

To examine these patterns in our data we turned our attention to locations for which we have relatively dense sampling: Southeast Farallon Island, off the coast of California; Floreana Island, in the Galapagos; Gough Island, a remote island in the south Atlantic; the grounds of a schoolhouse in Centreville, Maryland; and horse stables in the towns of Laurel, North Potomac and Chevy Chase, Maryland. Overall 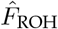 varies between populations (**Figure 7A**) (ANOVA: *F*_6,150_ = 45.5, *p <* 10^−6^). All individuals on Floreana and Farallon have high 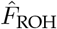 values. In Maryland, individuals range from effectively outbred (Centreville) to those that are as inbred as the mice on the small Pacific islands (Chevy Chase). Inbreeding on Gough Island, despite its remoteness, is relatively low. The empirical cumulative distributions of ROH segment lengths by individual are shown in (**Figure 7B**); there is clear difference in distribution between locations (*p <* 10^−6^, Anderson-Darling test), although this comparison does not take into account differences in total span of ROH. For purposes of qualitative comparisons we summarized ROH into bins of increasing size, from 2 cM (the detection threshold applied in this study) to *>* 20 cM (**Figure 7C**). To test for difference between geographic locations in aggregate ROH across length bins we used PERMANOVA, a non-parametric analogue to ANOVA for multivariate data. Binned ROH distributions do differ by population (pseudo-*F*_6,149_ = 36.8, permutation *p <* 10^−3^). Borrowing from analytical results and simulations in Ringbauer *et al*. (2021), it can be shown that segments 4 − 8 cM in length reflect background relatedness due to small population sizes while segments *>* 20 cM in length are only observed in matings between close kin or in very small populations (*N <* 500). Here a striking pattern emerges. Although inbreeding is high on the islands of Floreana (median 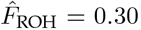) and Farallon (0.36), nearly all ROH lies in segments *<* 8 cM. The same is true on Gough Island, with the exception of a single individual. By contrast, in commensal populations in Maryland stables with similar overall inbreeding (Laurel, median 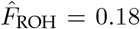; North Potomac, 0.26; Chevy Chase, 0.35) the majority of ROH lies in segments *>* 12 or *>* 20 cM. Inbreeding in the Centreville population is low (median 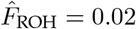).

**Figure 7:**
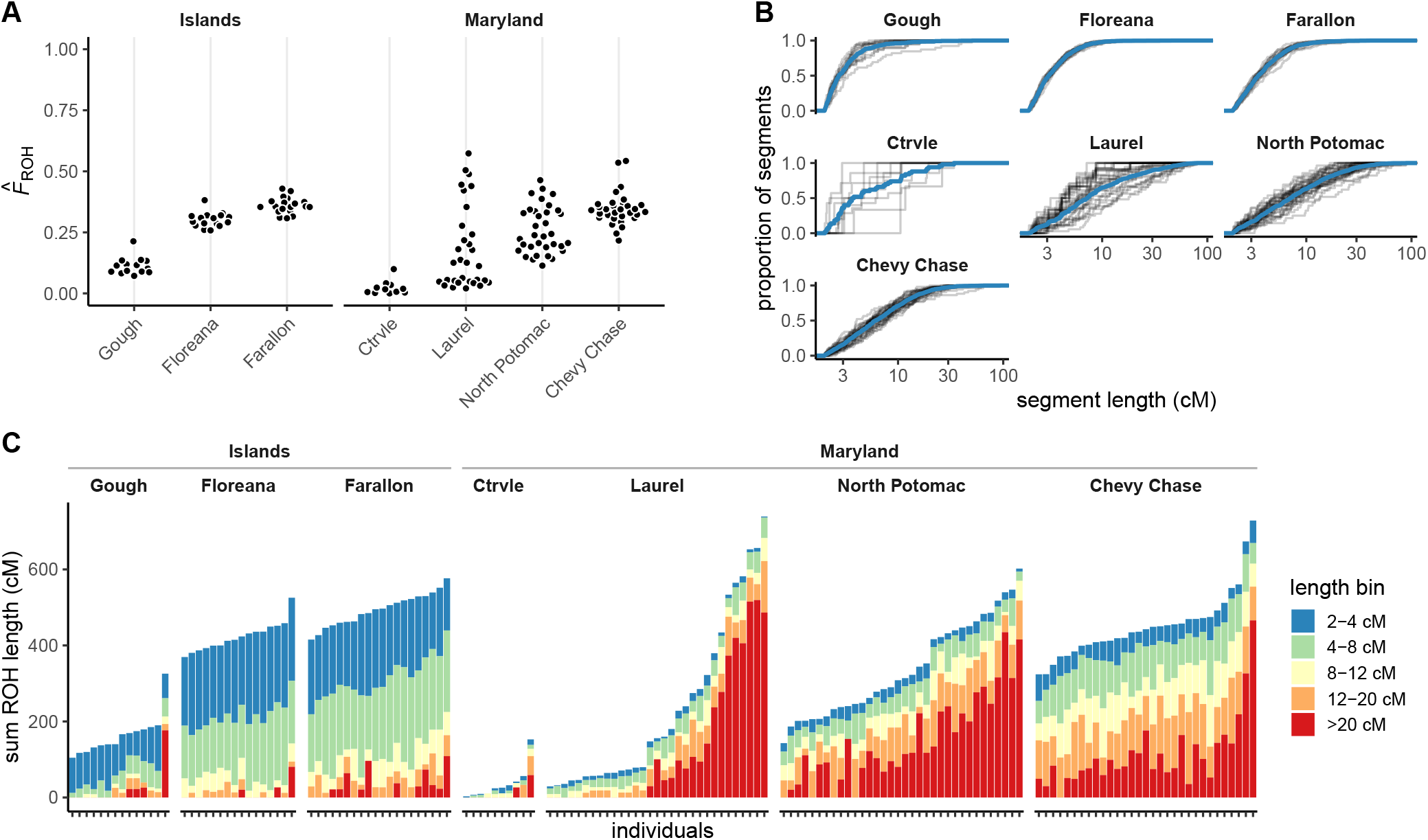
Inbreeding in selected deeply-sampled groups of *M. m. domesticus*. (**A**) Inbreeding coefficients 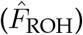 grouped by population for selected populations. Populations are sorted by mean 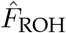. “Ctrvle” = Centreville, MD. (**B**) Cumulative distribution of ROH segment lengths (in cM) for each individual, grouped by population. Blue line shows pooled cumulative distribution for population. (**C**) Distribution of ROH by individual, binned by ROH length.

These results suggest that inbreeding in the Maryland populations is driven primarily by kin mating. We next examined patterns of relatedness within and between populations inferred from autosomal markers. The estimated kinship matrix is shown in **Figure 8A**, with rows and columns hierarchically clustered to emphasize close relationships. Relatedness is clearly higher within than between geographically separate groups, leading to block structure in the kinship matrix that corresponds to our geographically-defined population labels (shown by coloured bars along the edges of the matrix). We detect substructure even within populations that occupy the same horse barn, consistent with older literature (Selander 1970; Pocock *et al*. 2005). The distribution of pairwise kinship coefficients (*K*_*ij*_) is shown in **Figure 8B**. Within each of the three populations from horse stables (North Potomac, Laurel, Chevy Chase) inferred kinship for most pairs is at the level expected for first cousins (0.0625) or closer, with many pairs as similar as full siblings (0.25). (It is important to note that these expected values assume unrelated parents.) The range of kinship coefficients as well as the proportion of pairs with *K*_*ij*_ ≈ 0 is largest in Laurel. This is also the population with the broadest range of 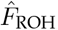, including 45% of individuals without ROH *>* 20 cM (see **Figure 7C**). The distribution of kinship coefficients in the Chevy Chase population is unimodal, centred at *K*_*ij*_ = 0.18, inbreeding is high (median 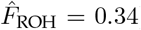) and all but one individual have two or more ROH *>* 20 cM in length. Mice in each barn are thus effectively a clan comprising one or a few extended families. The characteristics of North Potomac are intermediate between Laurel and Chevy Chase. Within the Centreville population, only a few pairs are closely related.

**Figure 8:**
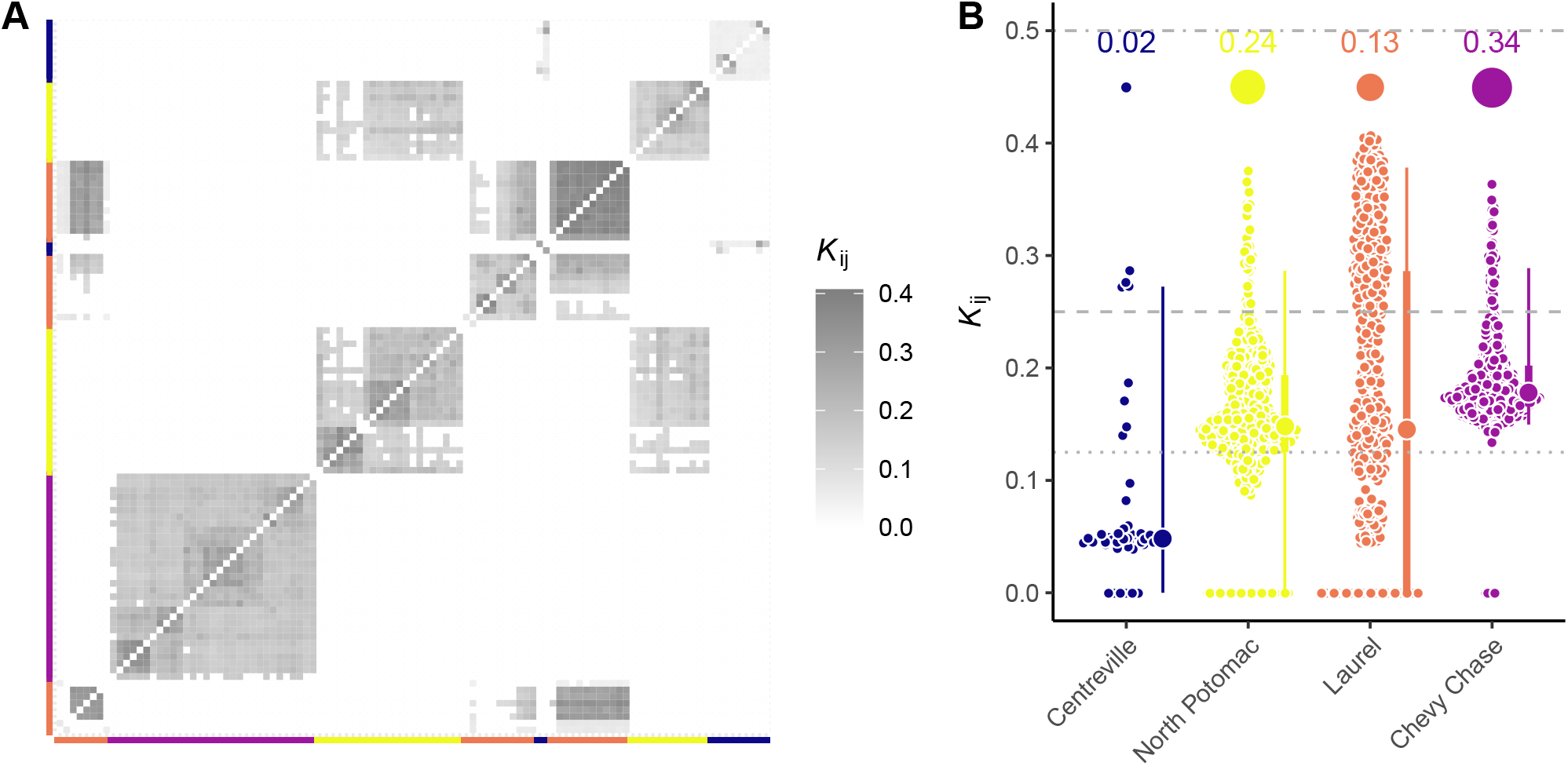
Deme structure in selected populations. (**A**) Kinship matrix (estimated from autosomal genotypes) for Maryland mice, hierarchically clustered. Row and column colours indicate population membership. (**B**) Distribution of pairwise kinship coefficients (*K*_*ij*_) within populations. Large dot, median; thick bar, interquartile range; thin bar, 5^th^ - 95^th^ percentiles. Reference lines show expected values for half-siblings (0.125), full siblings (0.250) and monozygotic twins (0.500) in the absence of inbreeding. Filled dots above each column indicate mean 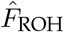 in each population.

## Discussion

The house mouse *Mus musculus* has been an important model organism in ecology, evolutionary biology, and medical science for more than a century. Here we present a survey of ancestry, population structure and inbreeding in a large and geographically diverse sample of wild mice (**Figure 1**) using genotypes at 57, 945 genome-wide SNP markers.

At a global scale, house mice comprise three major subspecies that are morphologically and genetically distinct and radiated from an ancestral population in central Asia (Boursot *et al*. 1993; Phifer-Rixey and Nachman 2015). *M. m. domesticus, M. m. musculus* and *M. m. castaneus* are well-differentiated in our data (**Figure 2**). We can detect substructure within *M. m. musculus* and *M. m. castaneus* but do not explore this in detail because of small sample size in these subspecies and ascertainment of SNPs primarily in *M. m. domesticus*. An additional group of mice collected in India, Pakistan, the Middle East, Madagascar and East Africa and collectively labelled “*M. musculus* undefined” in our study have genetic affinity with *M. m. castaneus*. Work by Hardouin *et al*. (2015), using nuclear microsatellites and with much more extensive sampling, showed clear evidence of *M. m. castaneus*-like ancestry in these regions as well as ancestry components distinct from any of the three major subspecies. This accords with previous evidence for deep branches within *M. m. castaneus* (Prager *et al*. 1998; Duplantier *et al*. 2002; Rajabi-Maham *et al*. 2012; Suzuki *et al*. 2013) based on mitochondrial sequences. Our study supports these findings but adds little additional detail. The taxonomic status of these populations remains uncertain.

We focus on population structure in *M. m. domesticus*, the dominant subspecies in western Europe and the Mediterranean. Through its commensal relationship with human colonists, this is also the subspecies that has expanded its range to the Americas, Australia, New Zealand and numerous islands around the globe within the past several centuries (Tichy *et al*. 1994; Searle *et al*. 2009a; Gabriel *et al*. 2010, 2011; Bonhomme and Searle 2012). Within the range of *M. m. domesticus* in Europe, we find clear evidence for a south-to-north ancestry gradient in multiple complementary analyses (**Figure 3,4,5**). Archaeological evidence and prior genetic surveys based primarily on mitochondrial sequences indicates that *M. m. domesticus* colonized southern and western Europe from the Levant via seafaring routes on the Mediterranean (Cucchi *et al*. 2005; Bonhomme *et al*. 2011; Bonhomme and Searle 2012; García-Rodríguez *et al*. 2018; Cucchi *et al*. 2020). This explains the southern grouping that we identify with our SNP data; the derivation and colonization routes of the central and northern European groupings that we have identified is less clear (**Figure 5B**). The colonization of central and northern Europe by *M. m. domesticus* from the Mediterranean could have occurred via overland or coastal routes (Cucchi *et al*. 2005; Jones *et al*. 2011). Previous studies suggest differences between the various defined mitochondrial DNA lineages in northern Europe in the manner of their arrivals (Jones *et al*. 2013). In the same way as we have here found a distinction between southern and northern Europe using genome-wide SNP data, the same can be seen with mitochondrial DNA: with inflated representation of particular lineages in the north which are not common in the south (Jones *et al*. 2011, 2013). Considering the next step of colonization, from Europe to elsewhere: mice in the eastern United States, from as far south as Florida to as far north as New Hampshire, have predominantly northern European-like ancestry and males carry a northern European Y haplogroup (**Figure 2B, 4A**). More specifically, mice from the eastern United States have ancestry profiles most similar to mice from New Zealand and Australia; and among mainland European populations, most similar to mice from mainland Scotland, Denmark and Germany (**Figure 4,S6**).

It is important to note that we have almost certainly not sampled the proximal source populations in Europe including, in particular, England and Scandinavia. Nonetheless, we can use populations we have sampled as imperfect proxies for the unsampled groups. Our data suggest that North American house mouse populations are likely descended primarily from populations in northern Europe. This is despite extensive Spanish activity in southern parts of North America from the sixteenth century CE onwards, which might have been expected to leave a mark in the mouse populations. (Our results are consistent with a prior RFLP analysis that identified polymorphisms shared between mice from Brittany and Florida (Tichy *et al*. 1994).) Given the size of the North American continent and the distance between early European settlements, multiple introductions from different European sources seem overwhelmingly more likely than a single introduction event. We also note that *M. m. castaneus* ancestry is also present in the western United States (Orth *et al*. 1998). Considering Australia and New Zealand, where house mice also appear to have northern European ancestry from our analysis, mitochondrial DNA data indicate mouse colonization from the British Isles, consistent with known human visitations during the eighteenth century CE and subsequent colonization of those areas (Searle *et al*. 2009a; Gabriel *et al*. 2011). It seems unlikely that indigenous people brought house mice to the Americas, Australia or New Zealand; if they did, those populations have been replaced by subsequent waves of colonization.

There are other instances where house mouse provenance, based on our SNP data, match well with known human history. Mice from Floreana Island in the Galápagos have Iberian-like ancestry (**Figure 3B**). The first recorded European landing on the Galápagos was by a Spanish expedition in 1535, and the islands were visited by whaling ships and explorers from England and the United States over the next three centuries. A fire set by a crewmember of the whaling ship *Essex* burned over all of Floreana in 1820 and may have reduced, if not eliminated, any mouse population present (Jackson 1993). Our results suggest that the present-day mouse population is either a remnant of a population introduced either by initial Spanish contact, or introduced from another source with Iberian-like ancestry (perhaps another island in the archipelago or the Ecuadorian mainland). Mice on Madeira and Porto Santo have clear affinity to coastal populations in Portugal. This diverges from previous work showing greater similarity of Madeiran mitochondrial haplotypes to those from northern Europe than from Portugal (Förster *et al*. 2009); but is consistent with the only other study of nuclear markers (Britton-Davidian *et al*. 2007). The small islands off the coast of New Zealand seem to have been colonized from a common source with New Zealand, as has been reported previously (Veale *et al*. 2018). One interesting result is that the mice from the Orkney Islands off the northern tip of Scotland show more affinity to mice from central Europe (Belgium, Northern France, Germany, Switzerland) than to other mice from northern Europe. Detailed studies of mouse mitochondrial sequences show a northern European affinity (Searle *et al*. 2009b). However, it is notable that another species of small mammal, the common vole *Microtus arvalis*, was also introduced to Orkney by people, and the source area was in the vicinity of the Low Countries (Martínková *et al*. 2013).

An important finding in our study is the degree of inbreeding in many house mouse populations surveyed. We exploit the length distribution of runs of homozygosity (ROH), which represent segments shared IBD between parents (Broman and Weber 1999), to make inferences both about individual inbreeding coefficients 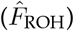 and population history (Kirin *et al*. 2010; Ceballos *et al*. 2018). ROH have several advantages for learning about recent population dynamics. First, ROH can be detected even with modest marker density and in the presence of marker ascertainment bias, and as such are a useful summary of genotyping array data from non-human and non-model species. Second, they contain information about population dynamics in the recent past (tens to a few hundred generations (Browning and Browning 2015), *<< N*_*e*_), whereas the site frequency spectrum is dominated by events deeper in time. These advantages extend beyond our study system. As might be expected, mice from many of the small islands surveyed (Floreana, Farallon, Heligoland, Gough, Orkney; **Figure 7**) are at the upper end of the distribution of 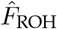 values. Historical surveys of other island populations based on allozyme data reached similar conclusions (reviewed in Berry 1986). However physical isolation, at least at a geographic scale, is neither necessary nor sufficient to account for observed levels of inbreeding. We took advantage of populations surveyed during a single season (autumn 2014) at several locations in Maryland to explore this in detail. Mice caught on the grounds of an elementary school in rural Centreville, MD are effectively outbred (**Figure 7**). By contrast, mice trapped in horse stables less than 100 km away, in an area with identical climate and very similar landscape, tend to be much more inbred. Homozygosity in these populations lies mostly in long segments (*>* 12 cM) that reflect recent consanguinity (**Figure 7**). Within stables, individuals are mutually related and can be lumped into large family groups within which there is a gradient of (realized) genetic relatedness (**Figure 8**).

These results support the idea that inbreeding in house mice is driven by fine-scale features of the habitat rather than broad geographic constraints. In a resource-rich environment like a horse barn – which may provide shelter, food and relative protection from predators – mice tend to live in small demes founded by up to a few dozen individuals with limited exchange between them, even over very short distances (Reimer and Petras 1967; Berry 1970; Lidicker 1976; Bronson 1979; Pocock *et al*. 2004). This of course leads to mating between close relatives, the signature of which is long segments of homozygosity. Mice in more austere habitats disperse over much larger distances and are less likely to encounter relatives among potential mates (Berry 1970; Bronson 1979; Pocock *et al*. 2005). In the case of isolated populations that have passed through a colonization bottleneck, such as on Farallon or Floreana, homozygosity may still be relatively high, but is distributed across shorter segments inherited identical by descent from the small pool of founder individuals (**Figure 7C**) (Ceballos *et al*. 2018).

Pervasive inbreeding seems to be at odds with estimates of nucleotide diversity in *M. m. domesticus*, which are about three-fold higher than in humans (for example Salcedo *et al*. 2007; Geraldes *et al*. 2008; Harr *et al*. 2016) despite similar mutation rates per generation. However, provided demes are sufficiently fluid over time, populations behave as if they are approximately panmictic at the regional scale, despite being finely subdivided on the local scale (Nagylaki 1977; Berry 1986). Furthermore, our data show that census population size need not be small – on Southeast Farallon Island, for example, may be very large (San Francisco Bay National Wildlife Refuge Complex 2013) – for levels of homozygosity to be high. This implies limitations on conclusions that can be drawn about population demographic history from genetic data alone. Users of genetics in conservation should be aware of these limitations.

Many open questions remain. How does inbreeding affect the genetic load and rate of adaptation in natural mouse populations? The fact that inbred strains can be generated readily in the laboratory may suggest that much deleterious recessive variation has been purged. While inbreeding is common in *M. m. domesticus*, is it equally common in *M. m. musculus, M. m. castaneus* and sister species *M. spretus*? And how does inbreeding affect the evolution of hybrid incompatibilities? High-quality whole genome sequences from a larger and more diverse panel of mice would be helpful to address these and other questions.

## Acknowledgements

This work was supported by grants from the following federal agencies: DARPA: D16-59 (DWT); National Institutes of Health: F30MH103925 (APM), T32GM067553 (APM, JPD), U19AI100625 (FPMdV), U42OD010924 (FPMdV), U24HG010100 (Leonard McMillan, FPMdV), P50GM076468 (Gary A Churchill, FPMdV); Oliver Smithies Investigator Award (FPMdV). JJH was supported by funding from the Cornell Center for Vertebrate Genomics and Cornell College of Agriculture and Life Sciences. We thank our many collaborators and sample collectors without whose extensive fieldwork this study would not have been possible, in particular the staff of Island Conservation on Southeast Farallon and Floreana. We thank Gerry McChesney of the US Fish and Wildlife Service for assistance with permits for trapping on Southeast Farallon Island. We thank the following technical staff for assistance with sample preparation, shipping and SNP array processing: Tim Bell, Ryan Buus, Jason Spence, Justin Gooch, Pablo Hock and Darla Miller. We thank Leonard McMillan and his research group for development of database systems to facilitate use of genotype data. Finally, we are grateful to Andrew Veale for making genotypes from New Zealand populations available to the community, and for discussions regarding their history.

## Data accessibility

Genotypes (**File S1**) and sample metadata (**Table S1**) have been deposited in Dryad at https://doi.org/10.5061/dryad.ncjsxkswt. New whole-genome sequence data have been deposited in the European Nucleotide Archive under accession number PRJEB52952.

## Author contributions

Contributed to study design and sample collection: APM, JJH, JPD, JBS, WJJ, KJC, DWT, FB, FPMdV. Analysed data: APM, JJH. Wrote paper: APM, JJH, JBS.

## Supplementary material

**Table S1**. Full sample information. See included column key for details.

**File S1**. Genotype matrix including all individuals (*N* = 814).

**Figure S1:**
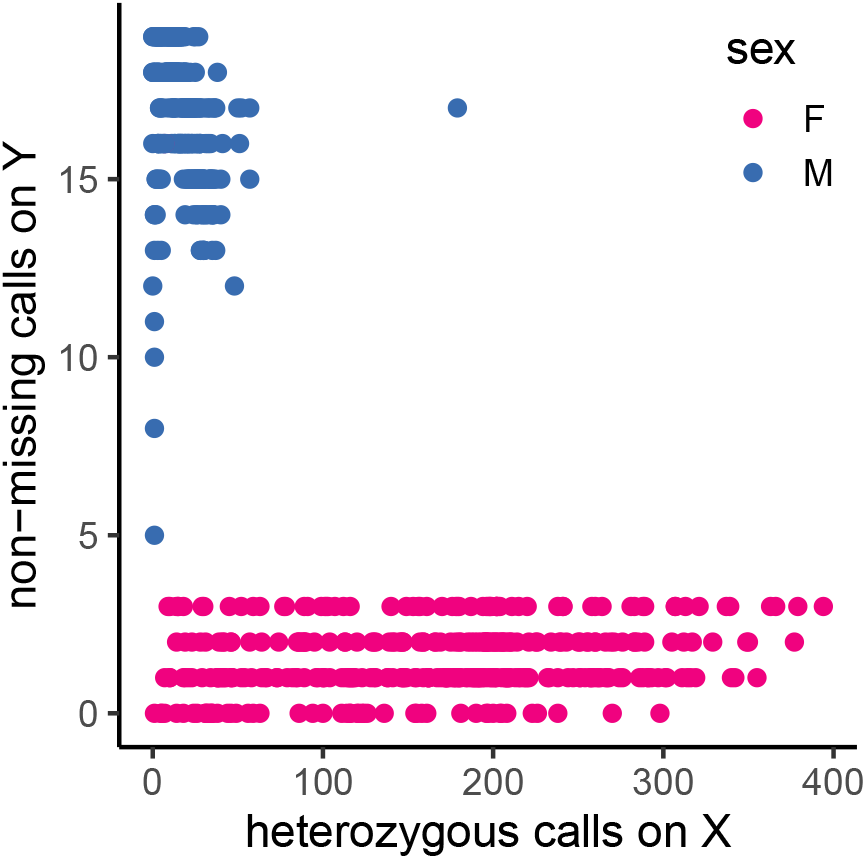
Confirmation of genetic sex. Each point represents one individual, colored by sex recorded at time of collection.

**Figure S2:**
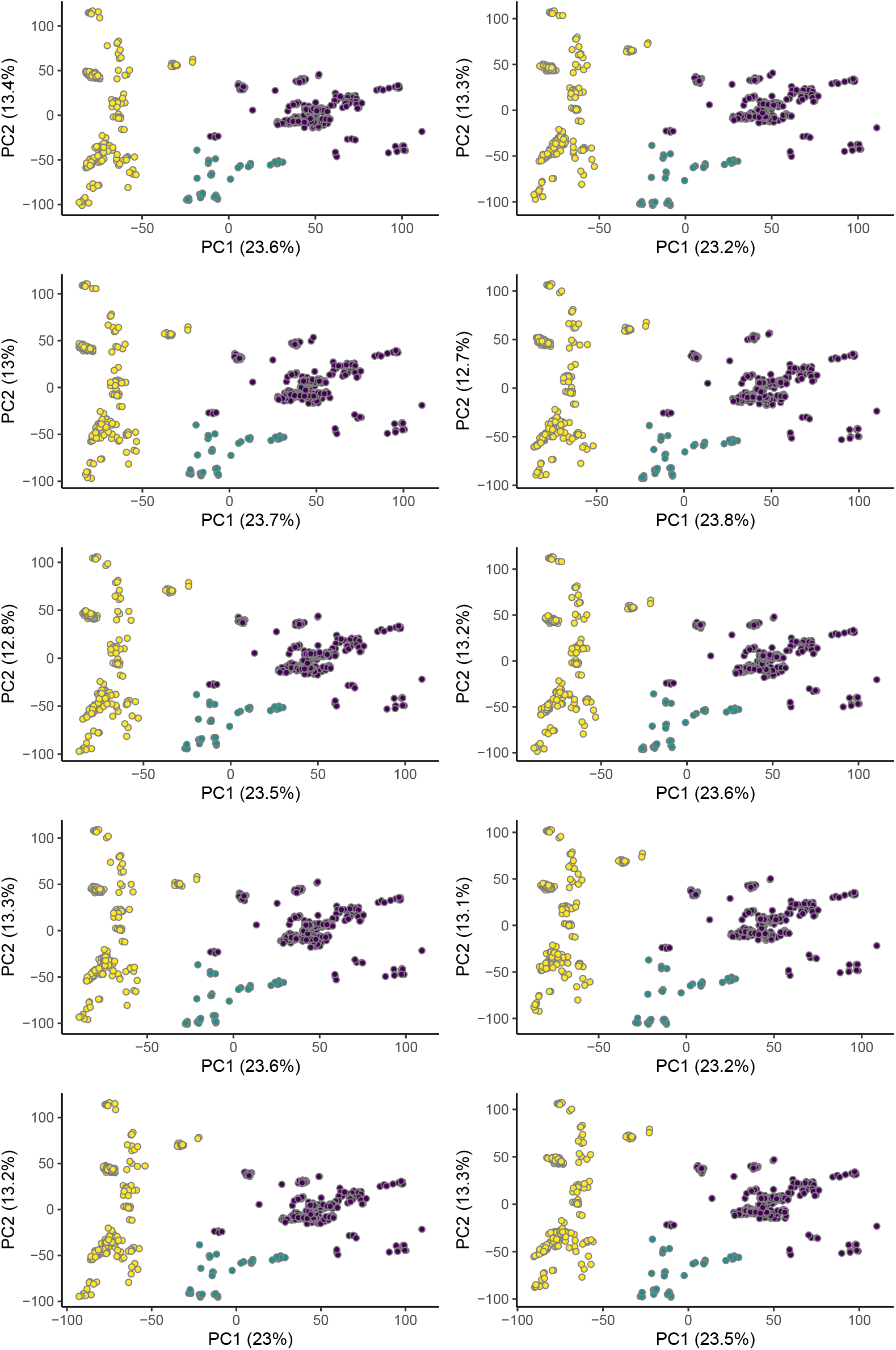
Population structure analyses are robust to choice of unrelated subset. Each panel shows PCA on autosomal genotypes using a different randomly sampled subset of unrelated *M. m. domesticus* individuals to calculate the principal components, and projecting the entire *M. m. domesticus* cohort onto these components. Color scheme follows **Figure 3A**.

**Figure S3:**
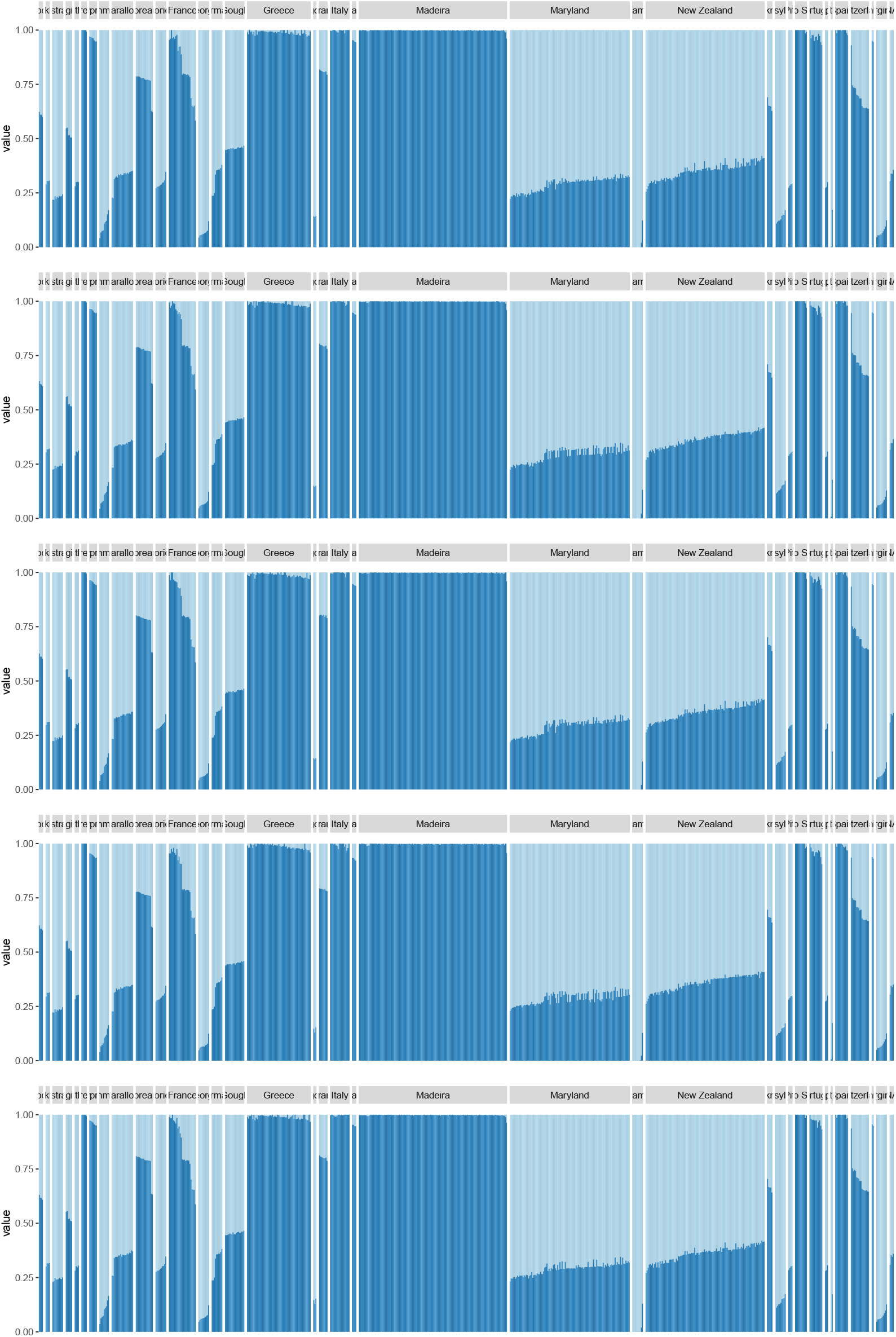
Ancestry decomposition is robust to choice of unrelated subset used to calculate allele frequencies. Each panel shows `ADMIXTURE` result using a different randomly sampled subset of unrelated *M. m. domesticus* individuals to estimate cluster-wise allele frequencies, then using these allele frequencies to perform ancestry decomposition on the entire cohort. Order of individuals is the same in each panel.

**Figure S4:**
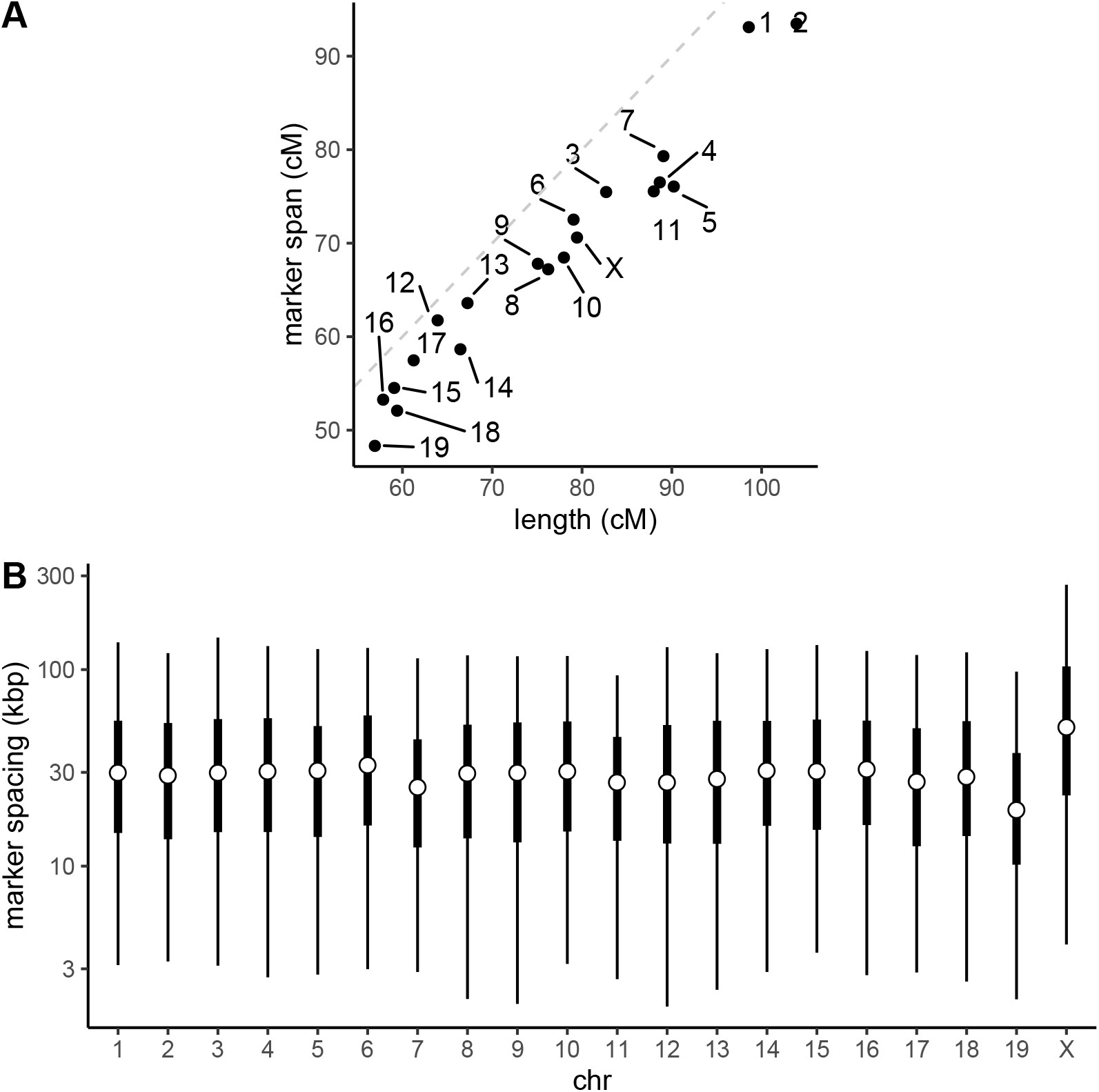
Genome coverage of marker panel used in this study. (**A**) Genetic span (in cM) of marker panel by chromosome versus estimated genetic length of each chromosome (Liu *et al*. 2014). (**B**) Physical spacing between adjacent markers by chromosome. Open dot, median; thick bar, interquartile range; thin bar, 5^th^ - 95^th^ percentiles.

**Figure S5:**
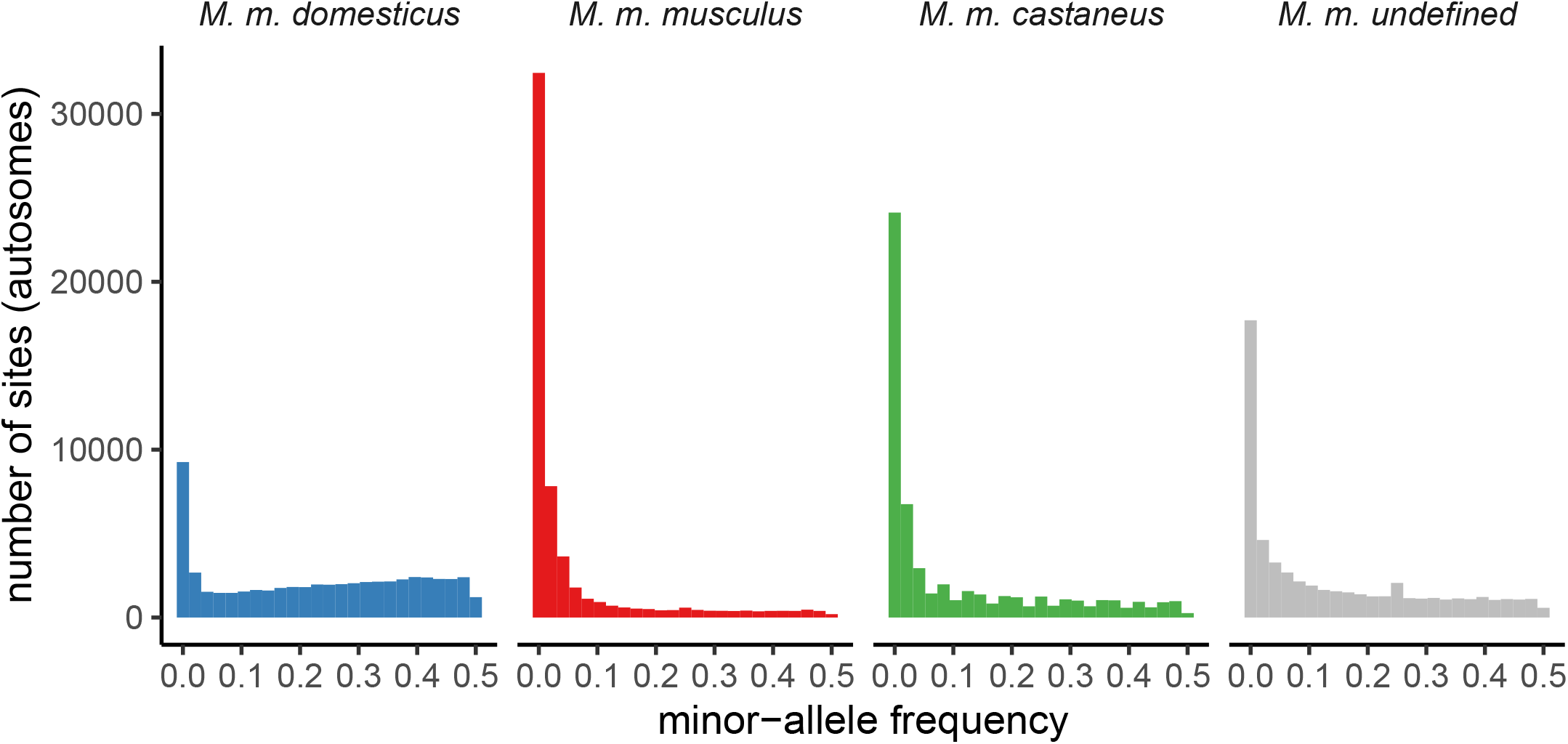
Folded site frequency spectrum by subspecies, for autosomal sites only.

**Figure S6:**
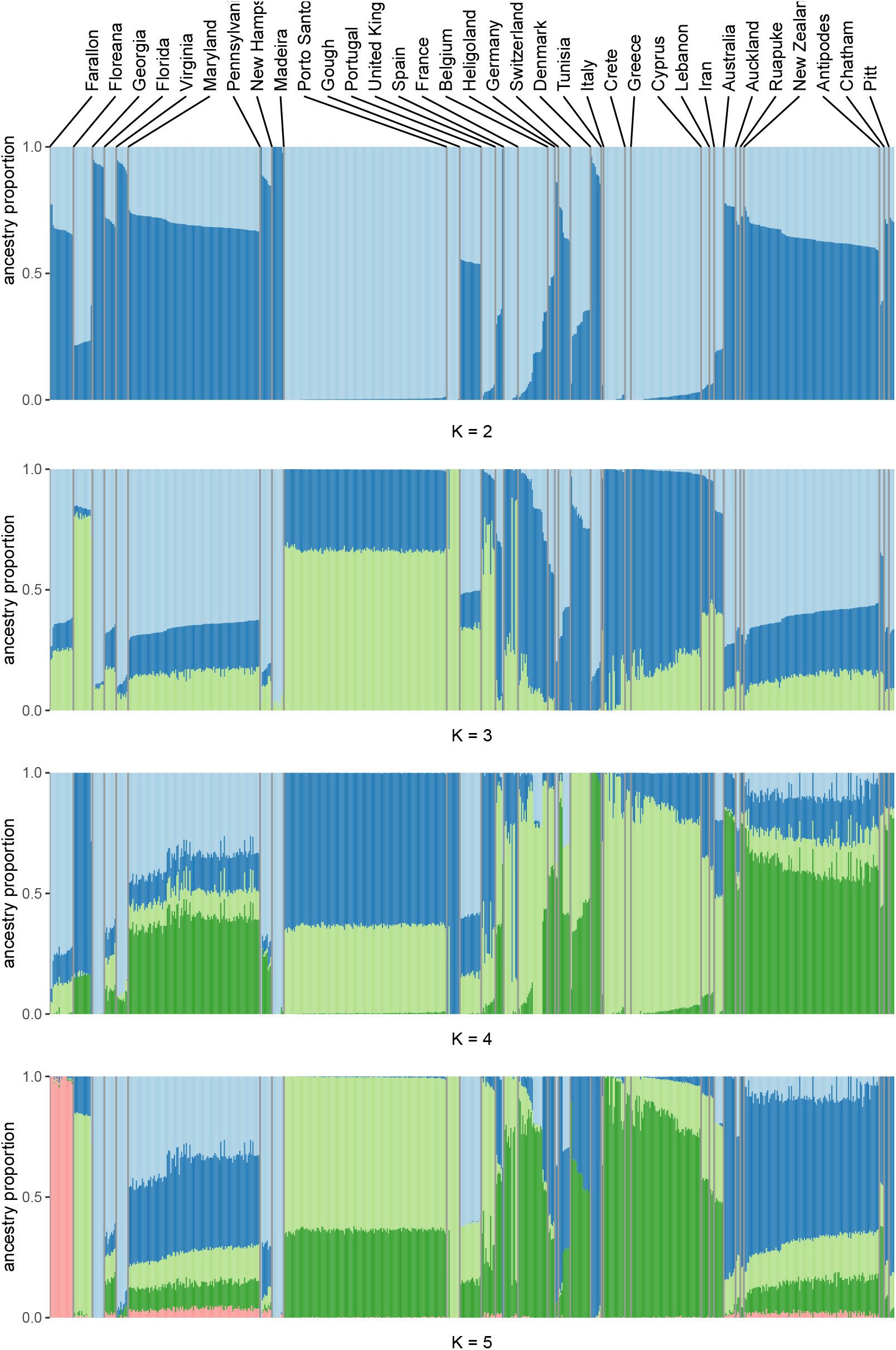
Ancestry proportions estimated with `ADMIXTURE` for *K* = 2, …, 5 mixture components. Populations sorted by longitude.

